# Brain Volumes, Thicknesses, and Surface Areas as Mediators of Genetic Factors and Childhood Adversity on Intelligence

**DOI:** 10.1101/2022.09.08.507068

**Authors:** Camille M. Williams, Hugo Peyre, Franck Ramus

## Abstract

Although genetic and environmental factors influence general intelligence (g-factor), few studies examined the neuroanatomical measures mediating environmental and genetic effects on intelligence. Therefore, we investigate the brain volumes, cortical mean thicknesses, and cortical surface areas mediating the effects of the g-factor polygenic score (gPGS) and childhood adversity on the g-factor in the UK Biobank.

We first identified the global and regional brain measures for the mediation models that contribute to the g-factor independently of global brain size and regional associations. Most regions contributed to the g-factor through global brain size. Parieto-Frontal Integration Theory (P-FIT) regions were not more associated with the g-factor than non-PFIT regions. Adjusting for global brain size and regional associations, only a few regions, such as the right olfactory sulcus and the right mediodorsal thalamic nuclei volumes, the right orbital inferior frontal gyrus surface area, and the anterior cingulate gyrus mean thicknesses, predicted intelligence and were included in the mediation analyses.

We conducted mediation analyses on global measures, regional volumes, mean thicknesses, and surface areas, separately. Total brain volume mediated 7.04% of the gPGS’ effect on the g-factor and 2.50% of childhood adversity’s effect on the g-factor. In comparison, the fraction of the gPGS and childhood adversity’s effects mediated by individual regional volumes, surfaces, and mean thicknesses was 10-15 times smaller. Our findings suggest that genetic and environmental effects on general intelligence must be mediated to a larger extent by other structural brain properties.

**Significance Statement:** Genes and environmental factors, such as childhood adversity, influence our cognitive abilities via the brain. Our results show that genetic and environmental effects on intelligence are mediated to some extent by neuroanatomical properties. However, we find that global brain measures (e.g., total brain volume) are the largest mediators and that regional volumes, surfaces, and mean thicknesses only mediated a fraction of a person’s genetic predisposition to intelligence and childhood adversity’s effect on intelligence. This suggests that genetic and environmental effects on general intelligence are to a large extent mediated by other kinds of brain structural properties.

## 1. Introduction

The positive correlation in performance across cognitive tests can be reduced to a single dimension: the general intelligence factor (g-factor), which reflects a person’s general cognitive performance. Although several studies examined the genetic and neurological basis of intelligence separately, there are relatively few studies investigating how genetic, environmental, and neurological factors simultaneously influence intelligence due to the lack of sufficiently rich and large datasets (for review Deary et al., 2021). Thus, this paper aims to capitalize on the richness of the UK Biobank – a large-scale prospective study with neural, genetic, environmental, and behavioral data - and identify the neuroanatomical measures (e.g., brain volumes) mediating the effect of genetic and environmental factors on intelligence.

Intelligence is heritable, with genetic differences accounting for about 50% of the differences in intelligence (Haworth et al., 2010; Polderman et al., 2015). Genome-Wide Association Studies (GWASs) identify genetic differences linked to variations in intelligence by pinpointing the Single-Nucleotide Polymorphisms (SNPs) that contribute to differences in intelligence. SNPs that vary with intelligence scores are typically associated with brain-expressed genes (Johnson et al., 2016; Lee et al., 2018) that are linked to a range of neuronal classes and processes, such as synaptic and neuron differentiation (Hill et al., 2019). SNP variations are therefore thought to be associated with differences at the macroscopic cerebral level.

SNP variations related to a trait can be summarized into a single score: a polygenic score (PGS), which reflects an individual’s genetic predisposition to a given phenotype. A PGS is derived from the sum of the effect allele at each SNP that is weighted by the SNP’s effect on a trait (estimated in a GWAS). The PGS of Cognitive Performance (i.e., measured by a verbal numerical score in the UK Biobank and a g-factor in Cognitive Genomics and Cohorts for Heart and Aging Research in Genomic Epidemiology consortiums) predicted up to 10.6% of the variance in cognitive performance in an independent sample (Lee et al., 2018). Although PGSs do not currently explain enough variance in intelligence to accurately predict individual intelligence scores (Morris et al., 2020), PGSs are valuable measures of genetic factors at the population level. Educational attainment and cognitive performance PGSs are increasingly used to disentangle environmental from genetic effects on educational and life outcomes (e.g., Bates et al., 2018; Rimfeld et al., 2018; Saarentaus et al., 2021; Stumm et al., 2020).

Since genetic and environmental effects act on intelligence via the brain, numerous studies investigated the neural correlates of intelligence. The most well-replicated association is the positive correlation of Total Brain Volume (TBV) with intelligence scores, ranging from r = 0.24 to 0.31 (Cox et al., 2019; Gignac & Bates, 2017; Pietschnig et al., 2015). Beyond overall brain size, the Parieto-Frontal Integration Theory (P-FIT; Jung & Haier, 2007) is the most supported theory on the regional correlates of intelligence. Although studies report additional regions than those predicted by the P-FIT, structural (grey and white matter volumes), diffusion, and functional studies find that intelligence scores are associated with the lateral and medial frontal, parietal, lateral temporal, and lateral occipital cortex, and their underlying white matter connectivity (e.g., arcuate fasciculus; e.g., Cox et al., 2019; Deary et al., 2021; Gur et al., 2021).

Previous UK Biobank studies examined the brain correlates of intelligence (Cox et al., 2016, 2019). The authors reported consistent associations with the P-FIT theory, such as stronger associations in the frontal pole, and the paracingulate, as well as less consistent associations with the P-FIT theory, such as weak associations in inferior frontal and superior parietal areas. They also found associations in the insula and precuneus/posterior cingulate volumes (Cox et al., 2019), which were more recently implicated in general intelligence (Basten et al., 2015). As for subcortical volumes, the UK Biobank study found that the thalamic volumes were the most associated with verbal numerical reasoning (β= 0.23). Finally, the authors reported that many of these regions still predicted intelligence when adjusting for TBV, suggesting that some regions make a unique contribution to intelligence that goes beyond TBV.

Although genetic and brain correlates of general intelligence (g-factor) have largely been studied, only two studies, to our knowledge, examined the extent to which neural measures mediate the effect of the g-factor polygenic score (gPGS) on the phenotypic g-factor. One study using vertex-wise mediation analyses of cortical thickness and cortical surface areas reported that the association between the gPGS and the phenotypic g-factor was mediated by the cortical thicknesses and surface areas of the anterior cingulate cortex, the prefrontal cortex, the insula, the medial temporal cortex, and inferior parietal cortex up to 0.75% in IMAGEN (N= 1,651) and 0.77% in IntegraMooDS (N= 742; Lett et al., 2020). In other words, these regions explained 20-40% of the variance explained by the gPGS on the g-factor (3-5%). A preprint on 550 adults, which used the same summary statistics of intelligence (Savage et al., 2018) as the above study to create their gPGSs, found that two intraparietal areas and the posterior temporal cortex surface areas mediated the effect of the gPGS on the g-factor (Genç et al., 2022). These mediation studies suggest that specific cortical regions mediate the effect of the gPGS on the phenotypic g-factor.

However, the extent to which additional regions, such as subcortical and cerebellar volumes, mediate the effect of the gPGS on intelligence has yet to be investigated. Moreover, the associations between cortical regions, genetics, and intelligence warrant further support and have not been investigated using the finer-grained segmentations of the Destrieux atlas. Therefore, our first aim was to examine whether subcortical volumes, cerebellar volumes, cortical volumes, cortical thicknesses, and cortical surface areas mediate the effects of the gPGS and the phenotypic g-factor with a more predictive gPGS in the UK Biobank.

Since early adversity is associated with a decrease in intelligence and cognitive function later in life (Enlow et al., 2012; McGuire & Jackson, 2020) and may have lasting biological and cerebral effects changes in childhood and adulthood (Dye, 2018; Lupien et al., 2009), our second aim was to examine whether the regions that mediate the gPGS’ effect on the g-factor also mediate childhood adversity’s effect on the g-factor. Taken together, this paper contributes to our understanding of the neuroanatomical measures mediating genetic (gPGS) and early environmental (childhood adversity) effects on intelligence (g-factor).

## 2. Methods

Analyses were run on R (R Core Team, 2022) and preregistered here: https://osf.io/ec97u/?view_only=4b366bd7ed2442a1a9f64bfcc2fe0946

### 2.1. Participants

The UK Biobank is a large prospective study with phenotypic, genotypic, and neuroimaging data from more than 500,000 participants. Participants were recruited between 2006 and 2010, from the vicinity of 22 assessment centers in England, Wales, and Scotland, with an age range for inclusion of 40–69 years. Data collection continues up to date.

All participants provided informed consent (“Resources tab” at https://biobank.ctsu.ox.ac.uk/crystal/field.cgi?id=200). The UK Biobank received ethical approval from the Research Ethics Committee (reference 11/NW/0382) and the present study was conducted based on application 46007.

We included participants whose combination of cognitive tests allowed for a correlation with the complete g-factor of 0.70 or higher (N= 261,701; Williams et al., 2022). This threshold was used to maximize the robustness of the factor and the number of participants with a g*-*factor.

In a previous paper, we analyzed the Image-Derived Phenotypes from the first Magnetic Imaging Resonance (MRI) visit generated by an image-processing pipeline developed and run by the UK Biobank Imaging team (Alfaro-Almagro et al., 2018; Miller et al., 2016) and reported 40,028 individuals with sex, age at MRI, and TBV data after excluding outliers (Williams et al., 2021). From here on, age at the first MRI visit will be referred to as age. From the 40,028 individuals with neuroimaging data, there were 39,131 participants with a g-factor of good quality (Table 1).

**Table 1.**
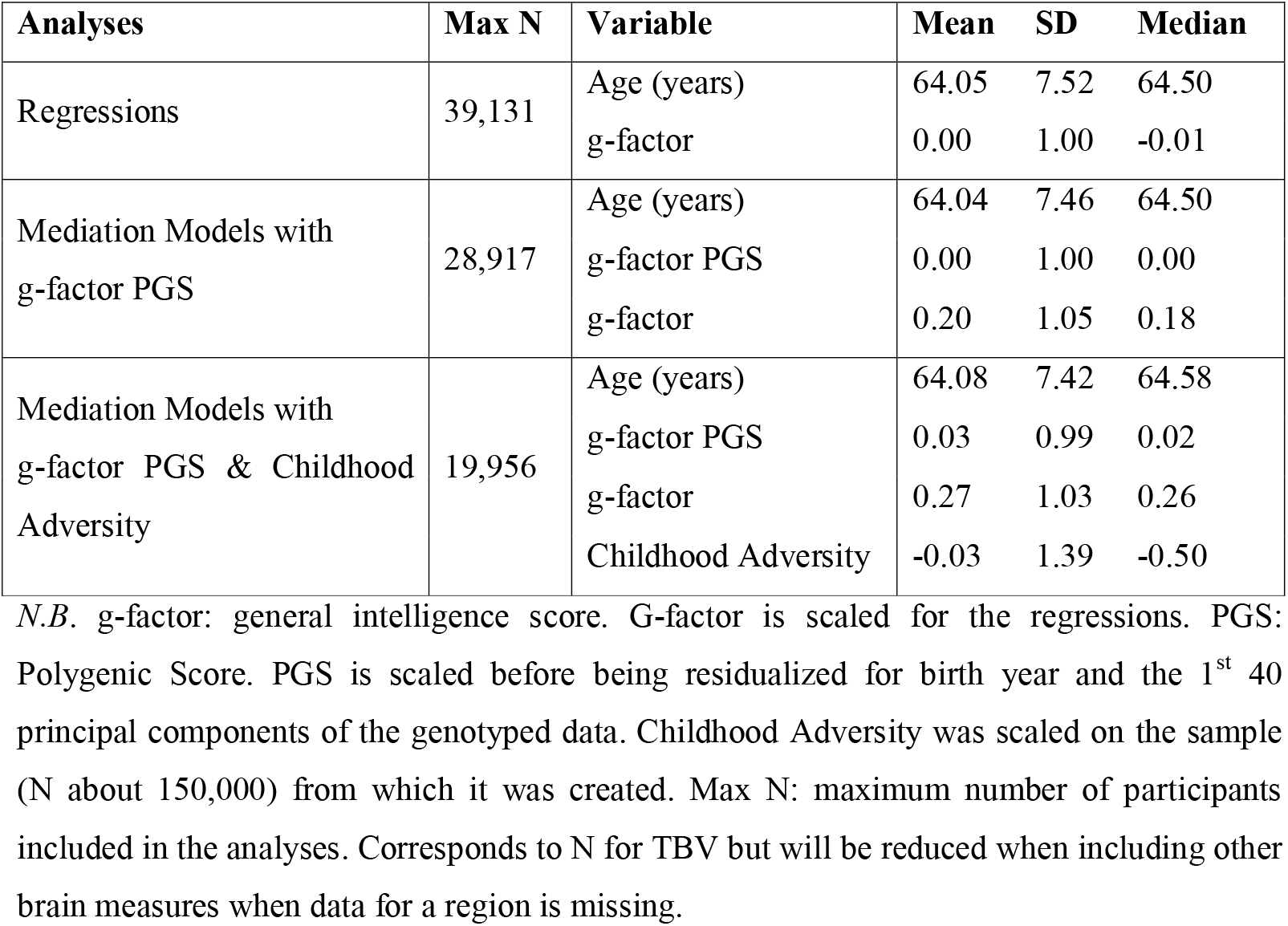
Descriptive Statistics of the Variables included in Each Analysis.

| **Analyses** | **Max N** | **Variable** | **Mean** | **SD** | **Median** |
| --- | --- | --- | --- | --- | --- |
| Regressions | 39,131 | Age (years) | 64.05 | 7.52 | 64.50 |
|  |  | g-factor | 0.00 | 1.00 | -0.01 |
| Mediation Models with g-factor PGS | 28,917 | Age (years) | 64.04 | 7.46 | 64.50 |
|  |  | g-factor PGS | 0.00 | 1.00 | 0.00 |
|  |  | g-factor | 0.20 | 1.05 | 0.18 |
| Mediation Models with g-factor PGS & Childhood Adversity | 19,956 | Age (years) | 64.08 | 7.42 | 64.58 |
|  |  | g-factor PGS | 0.03 | 0.99 | 0.02 |
|  |  | g-factor | 0.27 | 1.03 | 0.26 |
|  |  | Childhood Adversity | -0.03 | 1.39 | -0.50 |

*N.B.* g-factor: general intelligence score. G-factor is scaled for the regressions. PGS: Polygenic Score. PGS is scaled before being residualized for birth year and the 1<sup>st</sup> 40 principal components of the genotyped data. Childhood Adversity was scaled on the sample (N about 150,000) from which it was created. Max N: maximum number of participants included in the analyses. Corresponds to N for TBV but will be reduced when including other brain measures when data for a region is missing.

### 2.3. Imaged derived Phenotypes

We used the 10 global and 620 regional imaging phenotypes previously examined by Williams and colleagues (2021). The global phenotypes include TBV, total Mean Cortical Thickness (MCT), Total Surface Area (TSA), subcortical Grey Matter Volume (GMV), cortical GMV, cerebral White Matter Volume (WMV), cerebellar GMV, cerebellar WMV, the brainstem volume, and cerebral spinal fluid (CSF), whereas the regional phenotypes include: 444 cortical regions (148 volumes, 148 surface areas, and 148 cortical thicknesses) from the Freesurfer a2009s segmentations (Destrieux Atlas, data-field 197), 116 whole segmentations and subsegmentations of the amygdala, hippocampus, and thalamus and subsegmentations of the brainstem (Freesurfer subsegmentations, data-field 191), 28 cerebellum GMV segmentations from the FAST segmentations (data-field 1101), and 32 subcortical, white matter, and ventricle volumes from the Freesurfer ASEG segmentations (data-field 190). Freesurfer subcortical segmentations for the caudate, putamen, accumbens, and pallidum were used instead of the preregistered FIRST volumes, for segmentation consistency with the other subcortical and cortical volumes which were segmented from Freesurfer.

Although we did not preregister that we would examine the effects of the left and right measure of the whole thalamus, hippocampus, and amygdala because we focused the association of their subsegmentations with the g-factor, we ran these exploratory analyses to facilitate result comparison with previous studies and examine whether associations at the subcortical sub-segmentation level manifested at the global level.

### 2.4. Childhood Adversity Score

A childhood adversity score was created from questions in the UK Biobank on childhood abuse and social stressors. Childhood abuse was measured with data fields 20488 “When I was growing up… People in my family hit me so hard that it left me with bruises or marks”) and 20490 (“When I was growing up… Someone molested me (sexually)”) and childhood stressors were measured with data fields 20487 (“When I was growing up… I felt that someone in my family hated me”), 20489 (“When I was growing up… I felt loved”), and 20491 (“When I was growing up… There was someone to take me to the doctor if I needed it”). All questions are rated from 0 (Never True) to 4 (Very Often True). So that all indicators are in the same direction, we subtracted data field 20489 and data field 20491 responses from 4 (reverse coding). We conducted a PCA on the scores from these questions and extracted the first principal component (PC1) as our measure of childhood adversity, which captured 42% of the variance across questions (Table 2).

**Table 2.**
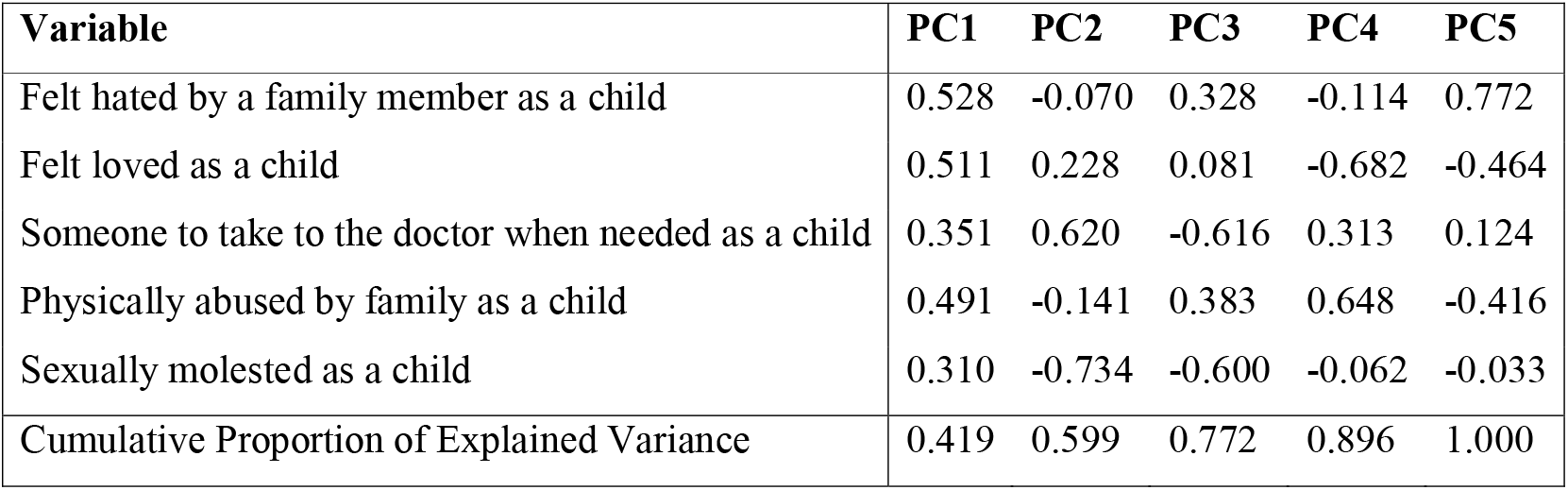
Principal Component (PC) Loadings of Childhood Abuse and Stressor Variables.

| Variable | PC1 | PC2 | PC3 | PC4 | PC5 |
| --- | --- | --- | --- | --- | --- |
| Felt hated by a family member as a child | 0.528 | -0.070 | 0.328 | -0.114 | 0.772 |
| Felt loved as a child | 0.511 | 0.228 | 0.081 | -0.682 | -0.464 |
| Someone to take to the doctor when needed as a child | 0.351 | 0.620 | -0.616 | 0.313 | 0.124 |
| Physically abused by family as a child | 0.491 | -0.141 | 0.383 | 0.648 | -0.416 |
| Sexually molested as a child | 0.310 | -0.734 | -0.600 | -0.062 | -0.033 |
| Cumulative Proportion of Explained Variance | 0.419 | 0.599 | 0.772 | 0.896 | 1.000 |

### 2.5. Statistical Analyses

We refer to the phenotypic g-factor as the g-factor and the g-factor PGS as the gPGS. A residualized gPGS was created by adjusting the gPGS for birth year and the first 40 principal components of the genotyped data and is referred to as the gPGS from here on out. All continuous variables were mean-centered and divided by 1SD. Females were coded 0.5 and Males – 0.5 in the regression analyses.

#### 2.5.1. What global measures predict the phenotypic g-factor?

We first estimated the effect of TBV and the CSF on the phenotypic g-factor, while adjusting for Sex, Age (quadratic and linear), their interactions, and Scanner Site (Equation 1, where i refers to an individual).

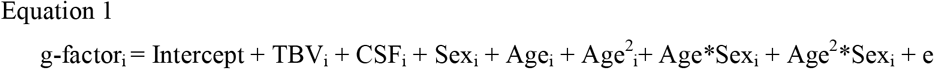

To identify the global measures driving the predictive effect of TBV on the g-factor, we simultaneously estimated the effect of total MCT, TSA, subcortical GMV cortical GMV, cerebral WMV, cerebellar GMV, cerebellar WMV, the brainstem volume, and CSF on the phenotypic g-factor, while adjusting for Sex, Age (quadratic and linear), their interactions, and Scanner Site (Equation 2, where i refers to an individual).

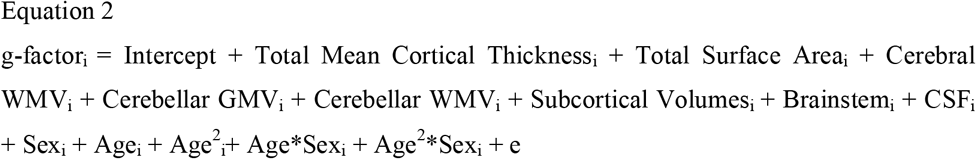

We considered that a regional measure significantly predicts the g-factor when p < 0.05/N, N: the number of coefficients of interest, which was 2 for equation 1 and 8 for equation 2.

#### 2.5.2. What global measures mediate the g-PGS’ effects on the g-factor?

We ran two mediation models using the sem function in the *lavaan* package (Rosseel, 2012): one with the significant predictors of the g-factor from equation 1 and one with those from equation 2. The gPGS was the exposure, global volume(s) the mediator(s), and the g-factor served as the outcome.

For all mediation models described in the present study, indirect effects were calculated using the product method. We estimated direct paths (exposure to an outcome) and indirect paths (exposure to mediators to outcomes) and adjusted the mediators and outcome for age (linear and quadratic), sex, and their interactions in the *lavaan* framework.

We set a lenient p-value threshold to 0.05 and a stricter one to p < 0.05/N (N: the number of regional and global measures included in the model of interest). Good fit was established with a CFI > 0.95, a RMSEA < 0.06 and a SRMR < 0.08 (Hu & Bentler, 1999).

#### 2.5.3. Do global measures mediate the g-PGS’ and Childhood Adversity’s effects on the g-factor?

We applied the same mediation models as in section 2.5.2. except that we added Childhood Adversity as additional exposure and estimated its direct and indirect paths through the global measure(s) to the g-factor. *Lavaan* considers correlations between predictors without estimating them.

#### 2.5.4. What regional measures predict the g-factor?

The aim was to identify the regions that contribute more to the g-factor than what is predicted given their size: this includes (1) regions that significantly predict the g-factor after adjusting for brain size and are positive with or without adjusting for global brain size and (2) regions that significantly predict the g-factor after adjusting for brain size and are negative with or without adjusting for global brain size.

To do so, we ran equations 3 and 4 (i refers to an individual, the regional measure N corresponds to a regional volume, thickness, or surface, and the global measure to TBV for volumes, Total MCT for mean thicknesses, and TSA for surface areas). The significance threshold was set to p < 0.05/N (N: the number of coefficients of interest, which was 148 for surfaces, 148 for thicknesses, and 311 for volumes).

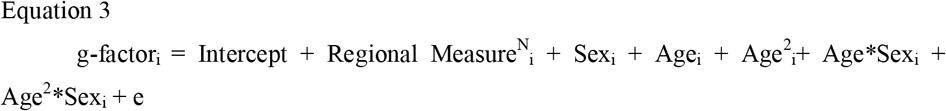

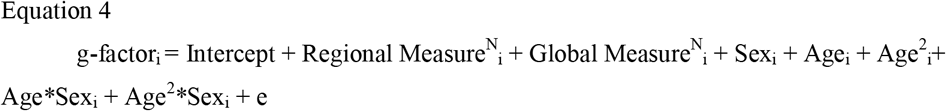

Although this was not preregistered, we tested whether the P-FIT theory accurately predicted the neuroanatomical measures most associated with the g factor. We mapped Brodmann’s Areas from the P-FIT (Colom et al., 2010; Haier & Jung, 2018; Jung & Haier, 2007) to the regions of the Destrieux Atlas based on the region names and the description of their location. The 14 Brodmann Areas that predict g according to the P-FIT were mapped to 62 of the 148 Destrieux segmentations. Several Destrieux regions were matched to the same Brodmann Area and several Brodmann Areas were matched to a single Destrieux region. Therefore, 12 Destrieux regions were matched twice to the Brodmann Areas, yielding 74 P-FIT Destrieux regions out of 160 Destrieux regions (Supplemental Table F1).

We then examined (1) whether P-FIT regions had a larger effect on the g-factor than non-P-FIT regions, by comparing the distribution of effect sizes of the two sets of regions with a t-test; and (2) whether P-FIT regions were overrepresented amongst the top 20 or 30 regions with the largest associations with the g-factor, using a chi-square test. We ran these analyses twice, for raw regional volumes, surfaces, and thicknesses, and for those adjusted on global brain measures. For the analysis of unadjusted regional volumes and surface areas, given that they were all significantly associated with the g factor, we restricted the analysis to the regions showing the top N associations.

#### 2.5.5. Do the Regional Measures that predict the g-factor independently from brain size still predict the g-factor when entered in the same model?

Based on equations 3 and 4, we selected the regions that still significantly and positively or negatively predicted the g-factor after adjusting for brain size. Because brain regions are correlated, their effect on the g-factor may be shared across regions even if they are independent of global brain size. Therefore, to avoid redundancy, we examined whether these regions still predicted the g-factor when simultaneously entered into a regression model predicting the g-factor (Equation 5, where N refers to a region, i to an individual, and the global measure to TBV for volumes, Total MCT for mean thicknesses, and TSA for surface areas). The significance threshold was set to p < 0.05/N (N: the number of regional measures included in the model of interest).

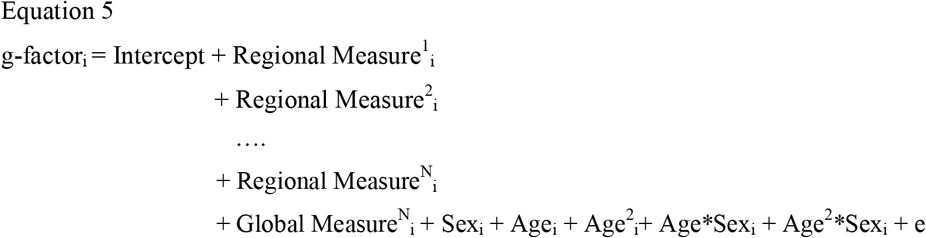

#### 2.5.6. Do Regional Measures mediate the g-PGS’ effects on the g-factor?

For volumes, thicknesses, and surface areas separately, we examined the simultaneous mediation of the global measure and the significant regional measures from equation 5 with the sem function from the *lavaan* package (Rosseel, 2012). The gPGS was the exposure, global and regional measures were the mediators, and the g-factor served as the outcome.

#### 2.5.7. Do Regional Measures mediate the g-PGS’ and Childhood Adversity’s effects on the g-factor?

We applied the same mediation models as in section 2.5.6. except that we added Childhood Adversity as additional exposure and estimated its direct and indirect paths through regional and global measures to g-factor.

## 3. Results

### 3.1. What global measures predict the phenotypic g-factor?

Greater TBV was associated with a greater g-factor (β = 0.24, SE = 0.006, p = 6.99e-297) and CSF did not predict the g-factor (Supplemental Table B1). When dividing TBV into its subcomponents, we found that greater TSA (β = 0.14, SE=0.012, p= 2.96e-31), Total MCT (β = 0.04, SE = 0.006, p = 7.11e-12), Cerebellar GMV (β = 0.08, SE = 0.008, p = 2.38e-26), and Cerebral WMV (β = 0.05, SE = 0.013, p = 1.03e-04) were associated with an increase in the g-factor (Figure 1; Supplemental Table B2). TBV explained 3.4% of the variance in the g-factor, whereas the global measures explained 3.6% of the variance in the g-factor

**Figure 1.**
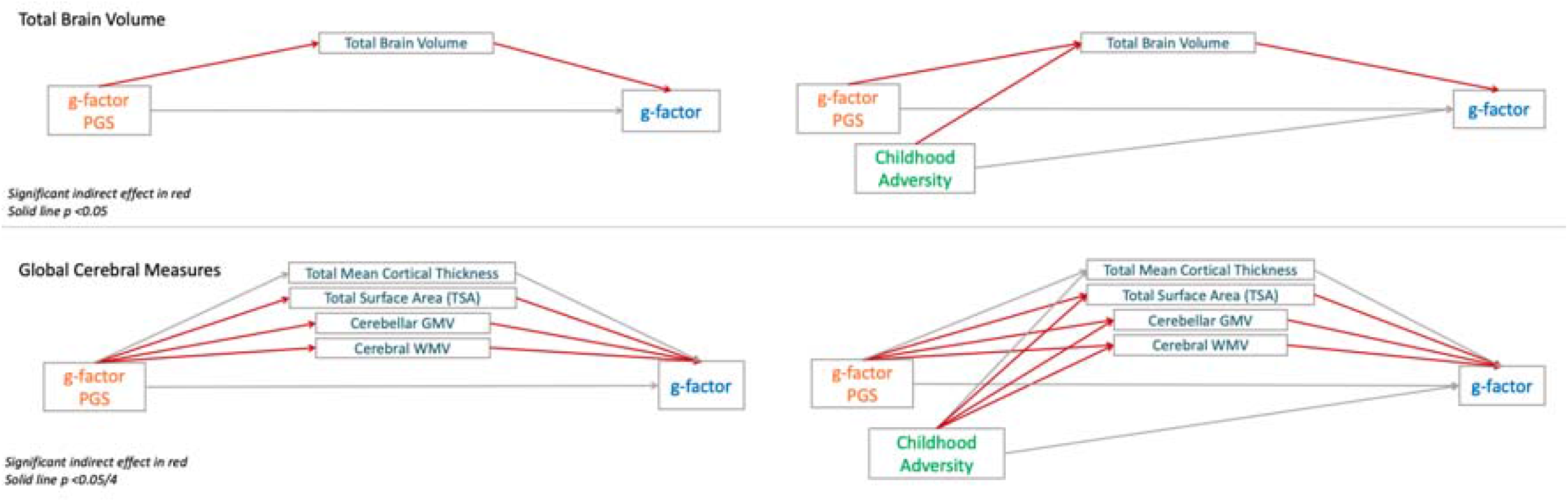
Meditating effect of global cerebral measures on the g-factor PGS’s effect on the g-factor with and without including Childhood Adversity. g-factor: general intelligence factor. PGS: polygenetic scores. Coefficients correspond to direct effects. Fit of TBV Models: CFI = 1.00, SRMR= 0.00, RMSEA = 0.00. Fit of Global Measures Model: CFI = 1.00, SRMR= 0.01, RMSEA = 0.03. The PGS is adjusted for birth year and the 1^st^ 40 principal components of the genotyped data. Cerebral measures and the g-factor are adjusted for sex, age, age^2^, age by sex, age^2^ by sex, and scanner site.

### 3.2. What global measures mediate the gPGS’ effect on the g-factor?

In the mediation model with TBV as the sole global mediator, TBV mediated 5.70% of the gPGS’ effect on the g-factor (Supplemental Table D1).

In the mediation model with several global measures as mediators, TSA mediates 3.32%, Cerebellar GMV 1.01%, and Cerebral WMV 1.28% of the gPGS’ effect on the g-factor (Figure 1; Supplemental Table D2).

### 3.3. What global measures mediate the gPGS and Childhood Adversity’s effect on the g-factor?

In the mediation model with TBV as the sole global mediator, TBV mediated 7.03% of the gPGS’ effect on the g-factor and mediated 2.49% of Childhood Adversity’s effect on the g-factor (Supplemental Table E1).

In the mediation model with several global measures as mediators, TSA mediates 3.68%, Cerebellar GMV 1.88% and Cerebral WMV 1.47% of the gPGS’ effect on the g-factor and TSA mediates 1.19%, Cerebellar GMV 0.56%, and Cerebral WMV 0.96% of Childhood Adversity’s effect on the g-factor (Figure 1; Supplemental Table E2).

### 3.4. What regional measures predict the g-factor?

Regression results are available in Supplemental Tables B (full models) and C (regional estimates). Figure 2 shows the g-factor estimate by volume, surface area, or thickness estimate when including or excluding the global measure in the regression model. Figures of the estimate by region for each type of possible change in significance or estimate (Table 3) are available in Files S1-3.

**Figure 2.**
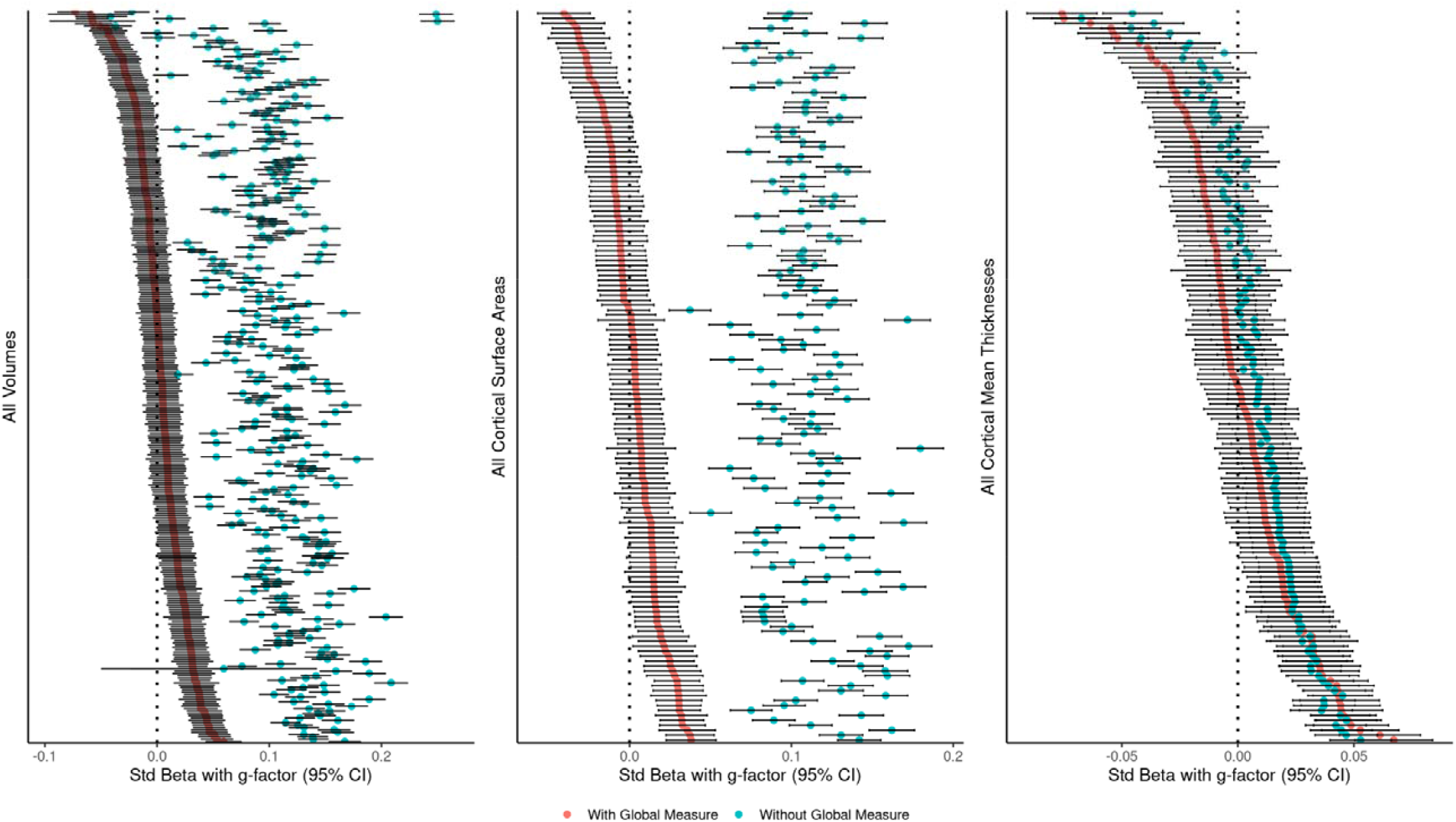
Standardized Estimate (Std Beta) of a Region’s association with the g-factor with (pink) and without (blue) adjusting for a Global Measure. Each point corresponds to a region. Region names are not shown for clarity. Global Measure: Total Brain Volume for volumes. Total Surface Area for surface areas. Total Mean Cortical Thickness for thicknesses.

**Table 3.** Number of Volumes, Surface Areas, and Mean Thicknesses by Types of Change in Significance and Estimate between Models without and with Global Brain Size.

| *Type of Change between Models* | **Volumes** |  | **Surface Areas** |  | **Mean Thicknesses** |  |
| --- | --- | --- | --- | --- | --- | --- |
|  | *N* | % | *N* | % | *N* | % |
| No Longer Significant | 242 | 77.81 | 130 | 87.84 | 6 | 4.05 |
| Still Significant Positive | 40 | 12.86 | 12 | 8.11 | 21 | 14.19 |
| Still Significant Negative | 2 | 0.64 | 0 | 0.00 | 7 | 4.73 |
| Still Not Significant | 5 | 1.61 | 0 | 0.00 | 104 | 70.27 |
| Still Significant, Positive to Negative | 18 | 5.79 | 6 | 4.05 | 0 | 0.00 |
| Becomes Significant & Negative | 4 | 1.29 | 0 | 0.00 | 10 | 6.76 |
| *Total* | 311 | 100.00 | 148 | 100.00 | 148 | 100.00 |

*N.B.* Global Brain size: Total Brain Volume for Volumes, Total Mean Cortical Thickness for thicknesses, and Total Surface Areas for surface areas

There were 242 out of 311 volumes, 130 out of 148 surface areas, and 6 out of 148 mean thicknesses, that were no longer significant after adjusting for global brain size (Table 3; Supplemental B3-4 & C1-3; Files S1-3). We found that the size of volumes and surface areas was significantly and positively correlated with the magnitude of its association with the g-factor without adjusting for global measures and that this association was no longer significant after adjusting for global brain size (File S4). Therefore, regional volumes and surfaces mainly contributed to the g-factor through global brain size.

There were 40 volumes (mainly cerebellar GMVs, subcortical thalamic and hippocampal nuclei, and a few cortical volumes), 12 cortical surface areas (mostly frontal), and 21 cortical mean thicknesses (mostly temporal; Table 3; Supplemental B5-6 & C1-3; Files S1-3) that were significant and had positive estimates after adjusting for brain size, suggesting that they contribute more (positively) to the g-factor than what is expected given their size.

There were 2 ventricular volumes and 7 cortical mean thicknesses (the left and right pericallosal sulci, the left and right anterior cingulate gyri and sulci, right occipital pole, left suborbital sulcus, and the right frontal marginal gyrus and sulcus; Table 3; Supplemental B5-6 & C1-3; Files S1-3) that were significant and had negative estimates after adjusting for brain size, suggesting that they contribute more (negative) to the g-factor than what is expected given their size.

There were 18 volumes and 6 surface areas that were still significant but had their estimates switch from positive to negative, suggesting that they contribute less to the g-factor than what is expected given their contribution to global brain size. For volumes, these regions included the right and left caudate, the right and left lingual gyrus, left and right pericallosal sulci, the right posterior-ventral part of the cingulate gyrus (isthmus), the right occipital pole, the right superior parietal gyrus, the hippocampal tail, and several subthalamic nuclei. As for surfaces, these regions included the left and right postcentral sulci, the left paracentral gyrus and sulcus, the middle anterior cingulate gyrus sand sulcus, the left lingual gyrus, and the left posterior-ventral part of the cingulate gyrus (isthmus).

Finally, there were 10 mean thicknesses, 3 ventricular volumes, and 1 subthalamic nucleus volume that became significant and negative after adjusting for brain size, suggesting that they contribute less to the g-factor than what is expected given their size. For mean thicknesses, regions include left and right transverse frontopolar gyri and sulci, the right lingual gyrus, right suborbital sulcus, the right superior frontal gyrus, the right cuneus gyrus, the right occipital superior gyrus, the right posterior-ventral part of the cingulate gyrus (isthmus), and left occipital pole and left posterior transverse collateral sulcus.

Across adjusted and unadjusted volumes, surface areas and thicknesses, and the different variants of our analyses, P-FIT regions were never overrepresented amongst the 20 or 30 regions that were the most associated with g (Files S5-6) and they did not show larger associations with g than non-PFIT regions overall (File S7).

### 3.5. Do the Regional Measures that predict the g-factor independently from brain size still predict the g-factor when entered in the same model?

We then examined whether regions that significantly and positively or negatively predicted the g-factor after adjusting for brain size do so independently from each other. We found that there were 4 Volumes, 3 Surface Areas, and 12 Mean Thicknesses that still significantly predicted the g-factor independently from each other and global brain size (Supplemental B9-11).

### 3.6. Do Regional Measures mediate the g-PGS’ effects on the g-factor?

#### 3.6.1. Volumes

Based on the previous analyses in section 3.5, we included the right olfactory bulb, the left subcallosal gyrus, and the Right Mediodorsal Medial Magnocellular Thalamic Nuclei volumes in the regional and global volumetric mediation model. We did not include the 3^rd^ ventricle volume even if it was significant because we do not expect ventricular volumes to mediate genetic effects on intelligence.

The indirect path from the gPGS to the g-factor was significant for TBV and the right olfactory bulb volume after multiple comparison corrections and for the left subcallosal gyrus volume at the p < 0.05 threshold. TBV mediated 5.07% of the gPGS’ effect on the g-factor, whereas the right olfactory bulb volume mediated 0.46% and left subcallosal gyrus volume 0.31% of the gPGS’ effect on the g-factor (Figure 3; Supplemental Table D3).

**Figure 3.**
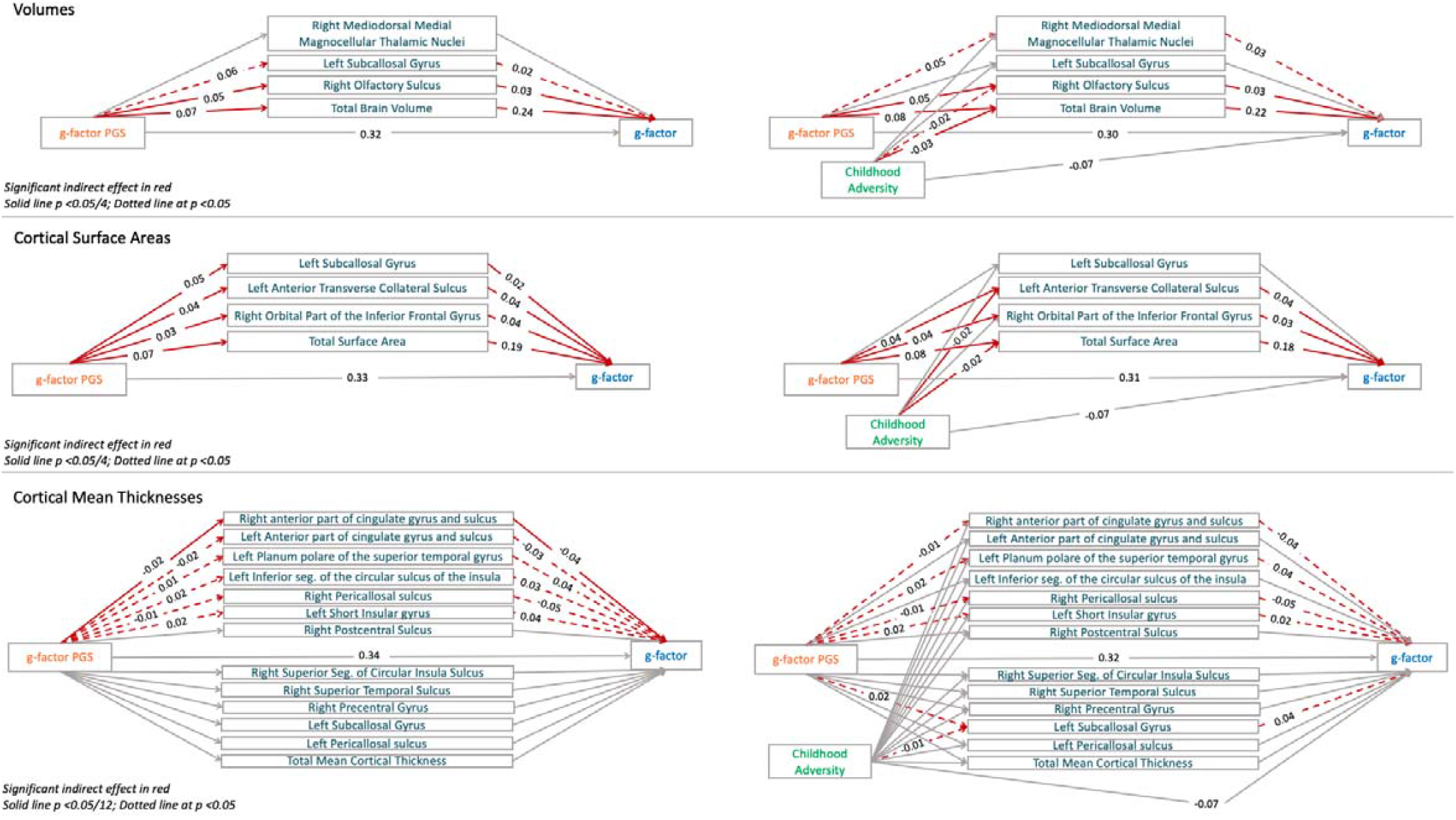
Meditating effect of regional cerebral measures on the g-factor PGS’s effect on the g-factor with and without including Childhood Adversity. g-factor: general intelligence factor. PGS: polygenetic scores. Coefficients correspond to direct effects. Fit of volume and surface area models: CFI = 1.00, SRMR= 0.00, RMSEA = 0.00. Fit of mean thickness model without Childhood Adversity: CFI = 0.95, SRMR= 0.05, RMSEA = 0.15, and with CFI = 0.95, SRMR= 0.05, RMSEA = 0.14. The PGS is adjusted for birth year and the 1^st^ 40 principal components of the genotyped data. Cerebral measures and the g-factor are adjusted for sex, age, age^2^, age by sex, age^2^ by sex, and scanner site.

#### 3.6.2. Surface Areas

Based on the previous analyses in section 3.5, we included TSA, the right orbital part of the inferior frontal gyrus surface area, the left subcallosal gyrus surface area, and the Left Anterior Transverse Collateral Sulcus surface area in the regional and global surface area mediation model.

Total Surface Area mediated 3.99%, the Left Anterior Transverse Collateral Sulcus surface area 0.41%, the left subcallosal gyrus 0.30%, and the Right Orbital Part of the Inferior Frontal Gyrus surface area 0.35% of the effect of the gPGS on the g-factor at the p < 0.05/4 threshold (Figure 3; Supplemental Table D4).

#### 3.6.3. Mean Thicknesses

Based on the previous analyses in section 3.5, we included Total MCT, the Right anterior part of the cingulate gyrus and sulcus, the left anterior part of the cingulate gyrus and sulcus, the left planum polare of the superior temporal gyrus, the left inferior segment of the circular sulcus of the insula, the right Pericallosal sulcus, the left Short Insular gyrus, the Right Postcentral Sulcus, the Right Superior Segment of Circular Insula Sulcus, the Right Superior Temporal Sulcus, the Right Precentral Gyrus, the Left Subcallosal Gyrus, and the Left Pericallosal sulcus mean thicknesses in the regional and global mean thickness mediation model.

Total MCT did not mediate the effects of the gPGS or Childhood Adversity on the g-factor. The right anterior part of the cingulate gyrus and sulcus mean thickness mediated 0.27% of the effect of the gPGS on the g-factor at p < 0.05. The left anterior part of the cingulate gyrus mediated 0.14%, the left short insular gyrus mean thickness 0.18%, the left planum polare of the superior temporal gyrus mean thickness 0.16%, the right pericallosal sulcus mean thickness mediated 0.12%, the left inferior segment of the circular sulcus of the insula mean thickness mediated 0.18%, and the right pericallosal sulcus mediated 0.27% of the effect of the gPGS on the g-factor at p < 0.05 (Figure 3; Supplemental Table D5).

### 3.7. Do Regional Measures mediate the g-PGS’ and Childhood Adversity’s effects on the g-factor?

#### 3.7.1. Volumes

In the volumetric mediation models, TBV mediated 4.15% of the gPGS’ effect on the g-factor and the right olfactory bulb volume mediated 0.37% of the gPGS’ effect on the g-factor at p < 0.05/4, whereas the right mediodorsal medial magnocellular thalamic nuclei volume meditated 0.32% of the gPGS’ effect on the g-factor at p < 0.05. Therefore, the left subcallosal gyrus volume was no longer significant when adding Childhood Adversity in the mediation model and the indirect path through the Right Mediodorsal Medial Magnocellular Thalamic Nuclei became significant at p < 0.05. TBV mediated 1.41% of Childhood Adversity’s effect on the g-factor, whereas the right olfactory bulb volume mediated 0.14% of Childhood Adversity’s effect on the g-factor at p < 0.05 (Figure 3; Supplemental Table E3).

#### 3.7.2. Surface Areas

Total Surface Area mediated 3.37%, the left anterior transverse Collateral Sulcus 0.34%, and the Right Orbital Part of the Inferior Frontal Gyrus surface area 0.29% of the effect of the gPGS on the g-factor at the p < 0.05/4 threshold (Supplemental Table D4). Total Surface Area mediated 1.11% and the Left Anterior Transverse Collateral Sulcus surface area 0.16% of Childhood Adversity’s effect on the g-factor (Figure 3; Supplemental Table E4).

#### 3.7.3. Mean Thicknesses

Total MCT did not mediate the effects of the gPGS or Childhood Adversity on the g-factor. The right anterior part of the cingulate gyrus and sulcus mean thickness still mediated 0.27% of the effect of the gPGS on the g-factor at p < 0.05. The right pericallosal sulcus mean thickness mediated 0.12%, the left short insular gyrus mean thickness 13%, and the left planum polare of the superior temporal gyrus mean thickness 0.15% of the effect of the gPGS on the g-factor at p < 0.05. However, the left inferior segment of the circular sulcus of the insula mean thickness and the left anterior part of the cingulate gyrus and sulcus mean thickness no longer mediated the gPGS’ effect on the g-factor. Instead, a new region mediated the gPGS’ effect on the g-factor: the left subcallosal gyrus mean thickness (0.17%), which also mediated the effect of Childhood Adversity’s effect on the g-factor (0.10%; Figure 3; Supplemental Table C5).

## Discussion

This paper capitalized on the richness of the UK Biobank to examine the extent to which neuroanatomical measures (e.g., brain volumes) mediate the effect of genetic and environmental factors on intelligence. We first examined the association between the g-factor and neuroanatomical measures in the UK Biobank on about 30,000 individuals with and without adjusting for global brain size and the remaining associations between regional measures. We then examined if global measures and regions that uniquely contribute to the g-factor mediate the gPGS’ effect on the g-factor and whether the same cerebral measures mediated childhood adversity’s effect on the g-factor. Although volumetric and surface area global measures were the main mediators of the gPGS’ effect on the g-factor, global measures mediated less than 10% of the gPGS’ effect on the g-factor and less than 3% of Childhood Adversity’s effect on the g-factor. Mediation by neuroanatomical measures, therefore, only explained a small fraction of the total effect of the gPGS and Childhood Adversity on the g-factor.

Most associations between the g-factor and volumes (78%) or surface areas (88%) disappeared after adjusting for global brain size, suggesting that the majority of volumes and surfaces contribute to intelligence through global cerebral effects. In contrast, only 4% of mean thicknesses were no longer significant after adjusting for global brain size and 26% of mean thicknesses still predicted the g-factor after adjusting for Total MCT. Therefore, mean thicknesses appear to influence the g-factor through region-specific effects rather than global effects. This can be explained by the small associations between regional mean thicknesses and Total MCT (mean β = 0.03) compared to those of regional volumes with TBV (mean β = 0.30) as well as the small association between Total MCT and the g-factor (β = 0.04). Adjusting for Total MCT thus captures little variance between regional mean cortical thicknesses and the g-factor.

Regions that contributed the most to the g-factor not only corresponded to regions that significantly predicted the g-factor after adjusting for brain size but also maintained the direction of their effect with and without adjusting for global measures. We found that cerebellar volumes, subcortical nuclei volumes, a few cortical volumes, frontal surface areas, and temporal mean thicknesses contributed more positively to the g-factor than what was expected given their size. In contrast, a few distributed mean thicknesses and ventricular volumes contributed more negatively to the g-factor than what was expected given their size. Negative and positive associations between cortical thickness and intelligence have been reported across the cortex (Karama et al., 2014) and are thought to depend on the measure of the intelligence (Goriounova & Mansvelder, 2019): Greater crystallized intelligence is associated with cortical thinning whereas fluid intelligence does not appear to be related with cortical thickness (Tadayon et al., 2020). Studies looking at age-related changes in performance on the cognitive tests of the UK Biobank found that performance on the verbal numerical UK Biobank test (also known as the fluid intelligence test) does not decrease as expected with age (Hagenaars et al., 2016). Instead, the performance stagnates as would be expected with crystallized intelligence (Cavanaugh & Blanchard-Fields, 2018). Therefore, our g-factor measure, on which the fluid intelligence test loads highly (0.62), likely captures both crystallized and fluid intelligence and may explain why we find distributed positive and negative associations between mean thicknesses and intelligence. Future studies should explore the associations between brain regions and subdomains of intelligence to obtain a better understanding of the associations between general intelligence, cognitive abilities, and the cortex (Jung & Haier, 2007).

When approximately matching the Destrieux segmentations to the P-FIT Brodmann Areas, the proportion of P-FIT and non-P-FIT regions associated with the g-factor was generally similar across cortical volumes, surface areas, and mean thicknesses and the magnitude of the association was not larger for P-FIT regions. Therefore, although we report associations with frontal and temporal-parietal regions that are concordant with the P-FIT, our findings do not support this theory with regard to volumes, thicknesses, and surface areas. However, the P-FIT may nonetheless accurately predict functional brain activations associated with general intelligence (Gur et al., 2021b; Haier & Jung, 2018).

Concerning other regions of interest, we did not replicate the significant associations between the g-factor and the whole hippocampal or thalamic volumes after adjusting for brain size previously reported by a UK Biobank study (Cox et al., 2019). Instead, we report associations between the g-factor and subcortical subsegmentations as well as cerebellar subsegmentations. For instance, we find a positive association between the g-factor and the right mediodorsal thalamic nucleus volume, a region known to critically contribute to cognitive functions (Ouhaz et al., 2018), and positive associations between the g-factor and most of the crus lobules of the cerebellum, which are functionally connected to regions of the default mode network (Buckner et al., 2011), a network of higher-level cognition (Smallwood et al., 2021).

We find that regional measures independently explained a small portion of the effect of the gPGS on the g-factor and that the gPGS’ effect on the g-factor is mediated by several cortical surface areas, volumes, and mean thickness. Although we used different segmentations and samples from Lett and colleagues (2020), we similarly find that the gPGS’ effect on the g-factor is mediated by the anterior cingulate cortex, prefrontal, insular, medial temporal, and inferior parietal mean thicknesses and surface areas. However, we find that our regional measures mediate a smaller percentage of the gPGS’ effect on the g-factor (around 0.30% instead of 0.75%), which may be due to the difference in age between cohorts or to their larger segmentations. We additionally report volumes that mediate the gPGS’ effect, and most notably the Right Mediodorsal Medial Magnocellular Thalamic Nuclei, a region thought to be implicated in executive functions (e.g., cognitive control and decision-making; Ouhaz et al., 2018).

When adding childhood adversity to the model, the percentage mediated by regional and global effects decreased to various extents across regions. For instance, adding childhood adversity to the mediation models did not impact the percentage mediated by the mean thickness of the right anterior part of the cingulate gyrus and sulcus. However, the surface area and volume of the left subcallosal gyrus and the mean thicknesses of the left inferior segment of the circular sulcus of the insula and the left anterior part of the cingulate gyrus and sulcus no longer mediated the gPGS’ effect on the g-factor when adding childhood adversity to the model. Considering that childhood adversity significantly and negatively predicted these regions, the association between these non-longer mediating regions may be due to the correlation between childhood adversity and the gPGS: Part of the variance previously attributed to the g-PGS may have shifted from the gPGS to childhood adversity. Finally, when adding childhood adversity to the mean thickness mediation model, a new region mediated both the gPGS effect and childhood adversity’s effect on the g-factor: the left subcallosal gyrus mean thickness. The latter highlights the importance of including environmental measures to better understand the complex relationship between environmental and genetic effects on general intelligence.

Although we find that specific regions mediate the g-factor and Childhood Adversity’s effects independently from global brain size and regional associations, the mediation of global brain size was 10-20 times larger than the mediation of specific regions when examining volumes and surface areas. TBV explained 2.3% and the regional volumes included in the mediation models explained 0.3% of the variance in the g-factor, whereas TSA explained 1.8% and the regional surface areas included in the mediation models explained 0.2% of the variance in the g-factor. These findings are consistent with previous studies suggesting that general intelligence may be more related to global than region-specific differences in the grey matter volume (Hilger et al., 2020) and that adding regional effects on the g-factor does not substantially predict more variance in the g-factor than TBV alone (Cox et al., 2019). However, TBV only mediated 7.04% of the gPGS’ effect on the g-factor and 2.50% of Childhood Adversity’s effect on the g-factor, leaving 93% of the gPGS’ effect on the g-factor, and 97.5% of Childhood Adversity’s effect on the g-factor to be explained by other cerebral measures. Therefore, future research should include additional cerebral measures, such as microstructural properties, white matter measures, dynamic connectivity, or resting-state or task-based functional activation, to better understand the extent to which cerebral measures mediate the environmental and genetic factors on the g-factor.

The present study is limited in its ability to generalize to all UK Biobank participants: Individuals with neuroimaging data are different from UK Biobank participants without neuroimaging data (Lyall et al., 2022) and the UK population (Fry et al., 2017). Moreover, our analyses were restrained to individuals of British ancestry, suggesting that further research is needed to examine whether genetic factors on intelligence are mediated by the same cerebral regions and to the same degree across ancestries. The polygenic score also only predicted 7.9% of the variance in the g-factor, suggesting that additional regions may be found when using a more predictive polygenic score. Finally, not including the most important environmental predictor of intelligence, parental or childhood socioeconomic status (SES; Flensborg-Madsen et al., 2020; Flensborg-Madsen & Mortensen, 2017), is a major limitation of our study. Since childhood SES was not available in the UK Biobank, we focused on childhood adversity, which is strongly associated with intelligence (but less so than parental SES; McGuire & Jackson, 2020). Although we could have used adult SES, as done by previous studies (e.g., Kweon et al., 2022) to serve as a proxy of childhood SES, the bidirectional influences between adult SES and intelligence would have compromised the interpretation of the mediation model.

The present paper provides the first large-scale study examining the neuroanatomical measures mediating genetic (gPGS) and early environmental (childhood adversity) effects on intelligence (g-factor). We replicate and extend previous findings and highlight the importance of adding environmental data to better understand the mechanisms by which genetic and environmental factors influence general intelligence. In light of the strong evidence for genetic and environmental factors contributing to individual differences in intelligence (for review Deary et al., 2021; Harden, 2021), we urge future studies to simultaneously investigate the genetic, environmental, and cerebral effects on intelligence by examining a variety of cerebral properties, from the macro to the micro, to understand discrepancies in intelligence (for review Deary et al., 2021), and, in turn, later health, educational, and social outcomes (Calvin et al., 2017; Schmidt & Hunter, 2004; Strenze, 2007; Twig et al., 2018).

## Acknowledgments

This work received support under the program “Investissements d’Avenir” launched by the French Government and implemented by l’Agence Nationale de la recherche (ANR) with the references ANR-17-EURE-0017 and ANR-10-IDEX-0001-02 PSL. A CC-BY public copyright license has been applied by the authors to the present document and will be applied to all subsequent versions up to the Author Accepted Manuscript arising from this submission, in accordance with the grant’s open access conditions. This research has been conducted using the UK Biobank Resource. Declarations of interest: none.

