## Supplementary material for "Brain Volumes, Thicknesses, and Surface Areas as Mediators of Genetic Factors and Childhood Adversity on Intelligence": File_S1_Types_of_Sig_Regions_Volumes.html

Volumes: Changes in Significance and Estimate Sign from Eq3 to Eq4


### Volumes: Changes in Significance and Estimate Sign from Eq3 to Eq4

### I - Table of number of regions by types of changes between Eq3 and Eq4

```
##                           Type of Change N Regions Percentage
## 1       Becomes Significant and negative         4       1.29
## 2                                Both NS         5       1.61
## 3              Both Significant Negative         2       0.64
## 4              Both Significant Positive        40      12.86
## 5 Both Significant, Positive to Negative        18       5.79
## 6                  No Longer Significant       242      77.81
```

### II - Plot of No Longer Significant Regions

There are too many regions to legibly include them in the plot.

### III - Plot of Still Significant Regions

#### A) Still Positive

#### B) Still Negative

### IV - Plot of Still Not Significant Regions

### V - Plot of Regions that become Significant and Negative

### VI - Plot of Regions that are still Significant but become Negative
