## Supplementary material for "Brain Volumes, Thicknesses, and Surface Areas as Mediators of Genetic Factors and Childhood Adversity on Intelligence": File_S2_Types_of_Sig_Regions_Surface_Areas.html

Surface Areas: Changes in Significance and Estimate Sign from Eq3 to Eq4


### Surface Areas: Changes in Significance and Estimate Sign from Eq3 to Eq4

### I - Table of number of regions by types of changes between Eq3 and Eq4

```
##                           Type of Change N Regions Percentage
## 1              Both Significant Positive        12       8.11
## 2 Both Significant, Positive to Negative         6       4.05
## 3                  No Longer Significant       130      87.84
```

### II - Both Significant

### III - No Longer Significant

There are too many regions to legibly include them in the plot.

### IV - Both Significant, Positive to Negative
