## Supplementary material for "Brain Volumes, Thicknesses, and Surface Areas as Mediators of Genetic Factors and Childhood Adversity on Intelligence": File_S3_Types_of_Sig_Regions_Mean_Thicknesses.html

Mean Thicknesses: Changes in Significance and Estimate Sign from Eq3 to Eq4


### Mean Thicknesses: Changes in Significance and Estimate Sign from Eq3 to Eq4

### I - Table of number of regions by types of changes between Eq3 and Eq4

```
##                     Type of Change N Regions Percentage
## 1 Becomes Significant and Negative        10       6.76
## 2                          Both NS       104      70.27
## 3        Both Significant Negative         7       4.73
## 4        Both Significant Positive        21      14.19
## 5            No Longer Significant         6       4.05
```

### I - Plots of Regions that are still not significant

There are too many regions to legibly include them in the plot.

### II - Plot of Still Significant Regions

#### A) Still Positive

#### B) Still Negative

### III - Plot of No Longer Significant Regions

There are too many regions to legibly include them in the plot.

### IV - Becomes Significant and Negative
