## Supplementary material for "Brain Volumes, Thicknesses, and Surface Areas as Mediators of Genetic Factors and Childhood Adversity on Intelligence": File_S4_Correlation_Regional_Size_and_Effect_Size.html

RStudio Sign In


Skip navigation


Error:

### Sign in to RStudio

Username:

Password:

Stay signed in when browser closes

You will automatically be signed out after 60 minutes of inactivity.


Sign In

### Signed Out

This browser was signed out from RStudio due to inactivity or by a manual sign out initiated from another tab. A new sign in was detected and you may now return to RStudio using the button below.

Return to RStudio
