## Supplementary material for "Brain Volumes, Thicknesses, and Surface Areas as Mediators of Genetic Factors and Childhood Adversity on Intelligence": File_S5_proportion_PFIT_nonPFIT_top_20.html

Cortical Associations with g by Theory with and without adjustment for global brain size


### Cortical Associations with g by Theory with and without adjustment for global brain size

Note. P-FIT : Parieto-frontal Integration Theory. 148: Number of cortical regions. N.S. : Not Significant. Global Brain size: Total Brain Volume for volumes, Total Surface Area for surface areas, and Total Mean Cortical Thickness for Mean Thicknesses.

### Proportion of PFIT regions among the top 20 associations.

#### 1. Cortical Surface Areas

##### a. Regression without Total Surface Area

```
##            
##             Not top 20 top 20
##   Not P-Fit        100      8
##   P-Fit             62     12
```

```
## 
##  Pearson's Chi-squared test with Yates' continuity correction
## 
## data:  con1
## X-squared = 2.6411, df = 1, p-value = 0.1041
```

```
## [1] "There were 12 P-FIT regions and 8 non P-FIT regions in the top 20 regions most associated with the g-factor. And there were 62  P-FIT regions and 100 non-PFIT regions that were not in the top 20."
```

```
## [1] "Therefore, P-FIT regions were as represented as non P-FIT regions in the top 20 regions associated with the g-factor (X2 (1, N = 160) = 2.641 p = 0.104."
```

##### b. Regression with Total Surface Area

```
##            
##             Not top 20 top 20
##   Not P-Fit         96     12
##   P-Fit             66      8
```

```
## 
##  Pearson's Chi-squared test with Yates' continuity correction
## 
## data:  con1
## X-squared = 0, df = 1, p-value = 1
```

```
## [1] "There were 8 P-FIT regions and 12 non P-FIT regions in the top 20 regions most associated with the g-factor. And there were 66  P-FIT regions and 96 non-PFIT regions that were not in the top 20."
```

```
## [1] "Therefore, P-FIT regions were as represented as non P-FIT regions in the top 20 regions associated with the g-factor (X2 (1, N = 160) = 0 p = 1."
```

#### 2. Mean Cortical Thickness

##### a. Regression without Total Mean Cortical Thickness

```
##            
##             Not top 20 top 20
##   Not P-Fit         90     18
##   P-Fit             72      2
```

```
## 
##  Pearson's Chi-squared test with Yates' continuity correction
## 
## data:  con1
## X-squared = 7.3845, df = 1, p-value = 0.006579
```

```
## [1] "There were 2 P-FIT regions and 18 non P-FIT regions in the top 20 regions most associated with the g-factor. And there were 72  P-FIT regions and 90 non-PFIT regions that were not in the top 20."
```

```
## [1] "Therefore, non P-FIT regions were more represented than P-FIT regions in the top 20 regions associated with the g-factor (X2 (1, N = 160) = 7.384 p = 0.007."
```

##### b. Regression with Total Mean Cortical Thickness

```
##            
##             Not top 20 top 20
##   Not P-Fit         91     17
##   P-Fit             71      3
```

```
## 
##  Pearson's Chi-squared test with Yates' continuity correction
## 
## data:  con1
## X-squared = 4.9949, df = 1, p-value = 0.02542
```

```
## [1] "There were 3 P-FIT regions and 17 non P-FIT regions in the top 20 regions most associated with the g-factor. And there were 71  P-FIT regions and 91 non-PFIT regions that were not in the top 20."
```

```
## [1] "Therefore, P-FIT regions were as represented as non P-FIT regions in the top 20 regions associated with the g-factor (X2 (1, N = 160) = 4.995 p = 0.025."
```

#### 3. Cortical Volumes

##### a. Regression without Total Brain Volume

```
##            
##             Not top 20 top 20
##   Not P-Fit         97     11
##   P-Fit             65      9
```

```
## 
##  Pearson's Chi-squared test with Yates' continuity correction
## 
## data:  con1
## X-squared = 0.031552, df = 1, p-value = 0.859
```

```
## [1] "There were 9 P-FIT regions and 11 non P-FIT regions in the top 20 regions most associated with the g-factor. And there were 65 P-FIT regions and 97 non-PFIT regions that were not in the top 20."
```

```
## [1] "Therefore, P-FIT regions were as represented as non P-FIT regions in the top 20 regions associated with the g-factor (X2 (1, N = 160) = 0.032 p = 0.859."
```

##### b. Regression with Total Brain Volume

```
##            
##             Not top 20 top 20
##   Not P-Fit         93     15
##   P-Fit             69      5
```

```
## 
##  Pearson's Chi-squared test with Yates' continuity correction
## 
## data:  con1
## X-squared = 1.6127, df = 1, p-value = 0.2041
```

```
## [1] "There were 5 P-FIT regions and 15 non P-FIT regions in the top 20 regions most associated with the g-factor. And there were 69  P-FIT regions and 93 non-PFIT regions that were not in the top 20."
```

```
## [1] "Therefore, non P-FIT regions were more represented than P-FIT regions in the top 20 regions associated with the g-factor (X2 (1, N = 160) = 1.613 p = 0.204."
```
