## Supplementary material for "Brain Volumes, Thicknesses, and Surface Areas as Mediators of Genetic Factors and Childhood Adversity on Intelligence": File_S6_proportion_PFIT_nonPFIT_top_30.html

Cortical Associations with g by Theory with and without adjustment for global brain size


### Cortical Associations with g by Theory with and without adjustment for global brain size

Note. P-FIT : Parieto-frontal Integration Theory. 148: Number of cortical regions. N.S. : Not Significant. Global Brain size: Total Brain Volume for volumes, Total Surface Area for surface areas, and Total Mean Cortical Thickness for Mean Thicknesses.

### Proportion of PFIT regions among the top 30 associations.

#### 1. Cortical Surface Areas

##### a. Regression without Total Surface Area

```
##            
##             Not Top 30 Top 30
##   Not P-Fit         94     14
##   P-Fit             58     16
```

```
## 
##  Pearson's Chi-squared test with Yates' continuity correction
## 
## data:  con1
## X-squared = 1.8038, df = 1, p-value = 0.1792
```

```
## [1] "There were 16 P-FIT regions and 14 non P-FIT regions in the top 30 regions most associated with the g-factor. And there were 58  P-FIT regions and 94 non-PFIT regions that were not in the top 30."
```

```
## [1] "Therefore, P-FIT regions were as represented as non P-FIT regions in the top 30 regions associated with the g-factor (X2 (1, N = 160) = 1.804 p = 0.179."
```

##### b. Regression with Total Surface Area

```
##            
##             Not Top 30 Top 30
##   Not P-Fit         89     19
##   P-Fit             63     11
```

```
## 
##  Pearson's Chi-squared test with Yates' continuity correction
## 
## data:  con1
## X-squared = 0.080549, df = 1, p-value = 0.7766
```

```
## [1] "There were 11 P-FIT regions and 19 non P-FIT regions in the top 30 regions most associated with the g-factor. And there were 63  P-FIT regions and 89 non-PFIT regions that were not in the top 30."
```

```
## [1] "Therefore, P-FIT regions were as represented as non P-FIT regions in the top 30 regions associated with the g-factor (X2 (1, N = 160) = 0.081 p = 0.777."
```

#### 2. Mean Cortical Thickness

##### a. Regression without Total Mean Cortical Thickness

```
##            
##             Not Top 30 Top 30
##   Not P-Fit         82     26
##   P-Fit             70      4
```

```
## 
##  Pearson's Chi-squared test with Yates' continuity correction
## 
## data:  con1
## X-squared = 9.8023, df = 1, p-value = 0.001743
```

```
## [1] "There were 4 P-FIT regions and 26 non P-FIT regions in the top 30 regions most associated with the g-factor. And there were 70  P-FIT regions and 82 non-PFIT regions that were not in the top 30."
```

```
## [1] "Therefore, non P-FIT regions were more represented than P-FIT regions in the top 30 regions associated with the g-factor (X2 (1, N = 160) = 9.802 p = 0.002."
```

##### b. Regression with Total Mean Cortical Thickness

```
##            
##             Not Top 30 Top 30
##   Not P-Fit         86     22
##   P-Fit             66      8
```

```
## 
##  Pearson's Chi-squared test with Yates' continuity correction
## 
## data:  con1
## X-squared = 2.2619, df = 1, p-value = 0.1326
```

```
## [1] "There were 8 P-FIT regions and 22 non P-FIT regions in the top 30 regions most associated with the g-factor. And there were 66  P-FIT regions and 86 non-PFIT regions that were not in the top 30."
```

```
## [1] "Therefore, P-FIT regions were as represented as non P-FIT regions in the top 30 regions associated with the g-factor (X2 (1, N = 160) = 2.262 p = 0.133."
```

#### 3. Cortical Volumes

##### a. Regression without Total Brain Volume

```
##            
##             Not Top 30 Top 30
##   Not P-Fit         93     15
##   P-Fit             59     15
```

```
## 
##  Pearson's Chi-squared test with Yates' continuity correction
## 
## data:  con1
## X-squared = 0.87676, df = 1, p-value = 0.3491
```

```
## [1] "There were 15 P-FIT regions and 15 non P-FIT regions in the top 30 regions most associated with the g-factor. And there were 59 P-FIT regions and 93 non-PFIT regions that were not in the top 30."
```

```
## [1] "Therefore, P-FIT regions were as represented as non P-FIT regions in the top 30 regions associated with the g-factor (X2 (1, N = 160) = 0.877 p = 0.349."
```

##### b. Regression with Total Brain Volume

```
##            
##             Not Top 30 Top 30
##   Not P-Fit         84     24
##   P-Fit             68      6
```

```
## 
##  Pearson's Chi-squared test with Yates' continuity correction
## 
## data:  con1
## X-squared = 5.3704, df = 1, p-value = 0.02048
```

```
## [1] "There were 6 P-FIT regions and 24 non P-FIT regions in the top 30 regions most associated with the g-factor. And there were 68  P-FIT regions and 84 non-PFIT regions that were not in the top 30."
```

```
## [1] "Therefore, non P-FIT regions were more represented than P-FIT regions in the top 30 regions associated with the g-factor (X2 (1, N = 160) = 5.37 p = 0.02."
```
