## Supplementary material for "Brain Volumes, Thicknesses, and Surface Areas as Mediators of Genetic Factors and Childhood Adversity on Intelligence": File_S7_effect_size_comparison_PFIT_nonPFIT.html

Cortical Associations with g by Theory with and without adjustment for global brain size


### Cortical Associations with g by Theory with and without adjustment for global brain size

Note. P-FIT : Parieto-frontal Integration Theory. 148: Number of cortical regions. N.S. : Not Significant. Global Brain size: Total Brain Volume for volumes, Total Surface Area for surface areas, and Total Mean Cortical Thickness for Mean Thicknesses. ABs. Absolute

### A) Comparison of effect sizes

Distribution of the ABSOLUTE standardized associations of each cortical region with the g-factor by Theory without adjustment for Global Brain Size.

We first ran an F-test to know whether to run a classic t-test (assumes equal variance across groups) or a Weltch t-test (does not assume equal variance across groups).

#### 1. All regions without global brain size adjustment

##### a. Volumes F-test, t-test

```
##      Theory  grp.mean
## 1 Not P-FIT 0.1042791
## 2     P-FIT 0.1058649
```

```
##      Theory     grp.sd
## 1 Not P-FIT 0.03114620
## 2     P-FIT 0.02885738
```

```
## 
##  F test to compare two variances
## 
## data:  Estimate_rounded by Theory
## F = 1.1649, num df = 85, denom df = 73, p-value = 0.5048
## alternative hypothesis: true ratio of variances is not equal to 1
## 95 percent confidence interval:
##  0.7432642 1.8119142
## sample estimates:
## ratio of variances 
##            1.16492
```

```
## 
##  Two Sample t-test
## 
## data:  Estimate_rounded by Theory
## t = -0.33215, df = 158, p-value = 0.7402
## alternative hypothesis: true difference in means is not equal to 0
## 95 percent confidence interval:
##  -0.011015496  0.007843906
## sample estimates:
## mean in group Not P-FIT     mean in group P-FIT 
##               0.1042791               0.1058649
```

##### b. Surface Areas F-test, t-test

```
##      Theory  grp.mean
## 1 Not P-FIT 0.1088023
## 2     P-FIT 0.1139189
```

```
##      Theory     grp.sd
## 1 Not P-FIT 0.02500391
## 2     P-FIT 0.02833931
```

```
## 
##  F test to compare two variances
## 
## data:  Estimate_rounded by Theory
## F = 0.77846, num df = 85, denom df = 73, p-value = 0.2653
## alternative hypothesis: true ratio of variances is not equal to 1
## 95 percent confidence interval:
##  0.4966888 1.2108177
## sample estimates:
## ratio of variances 
##           0.778462
```

```
## 
##  Two Sample t-test
## 
## data:  Estimate_rounded by Theory
## t = -1.2133, df = 158, p-value = 0.2268
## alternative hypothesis: true difference in means is not equal to 0
## 95 percent confidence interval:
##  -0.013446012  0.003212825
## sample estimates:
## mean in group Not P-FIT     mean in group P-FIT 
##               0.1088023               0.1139189
```

##### c. Mean Thicknesses F-test, t-test

```
##      Theory   grp.mean
## 1 Not P-FIT 0.02016279
## 2     P-FIT 0.01082432
```

```
##      Theory      grp.sd
## 1 Not P-FIT 0.014931893
## 2     P-FIT 0.009143241
```

```
## 
##  F test to compare two variances
## 
## data:  Estimate_rounded by Theory
## F = 2.667, num df = 85, denom df = 73, p-value = 2.714e-05
## alternative hypothesis: true ratio of variances is not equal to 1
## 95 percent confidence interval:
##  1.701674 4.148306
## sample estimates:
## ratio of variances 
##           2.667039
```

```
## 
##  Welch Two Sample t-test
## 
## data:  Estimate_rounded by Theory
## t = 4.8403, df = 143.49, p-value = 3.312e-06
## alternative hypothesis: true difference in means is not equal to 0
## 95 percent confidence interval:
##  0.005524899 0.013152033
## sample estimates:
## mean in group Not P-FIT     mean in group P-FIT 
##              0.02016279              0.01082432
```

##### d. Plot of Distribution

#### 2. All regions with global brain size adjustment

Distribution of standardized associations of each cortical region with the g-factor by Theory with adjustment for Global Brain Size.

##### a. Volumes F-test, t-test

```
##      Theory   grp.mean
## 1 Not P-FIT 0.01522093
## 2     P-FIT 0.01029730
```

```
##      Theory     grp.sd
## 1 Not P-FIT 0.01027893
## 2     P-FIT 0.00923028
```

```
## 
##  F test to compare two variances
## 
## data:  Estimate_rounded by Theory
## F = 1.2401, num df = 85, denom df = 73, p-value = 0.3465
## alternative hypothesis: true ratio of variances is not equal to 1
## 95 percent confidence interval:
##  0.7912493 1.9288914
## sample estimates:
## ratio of variances 
##           1.240128
```

```
## 
##  Welch Two Sample t-test
## 
## data:  Estimate_rounded by Theory
## t = 3.1916, df = 157.7, p-value = 0.001708
## alternative hypothesis: true difference in means is not equal to 0
## 95 percent confidence interval:
##  0.001876635 0.007970630
## sample estimates:
## mean in group Not P-FIT     mean in group P-FIT 
##              0.01522093              0.01029730
```

##### b. Surface Areas F-test, t-test

```
##      Theory   grp.mean
## 1 Not P-FIT 0.01463953
## 2     P-FIT 0.01201351
```

```
##      Theory      grp.sd
## 1 Not P-FIT 0.010057380
## 2     P-FIT 0.009872465
```

```
## 
##  F test to compare two variances
## 
## data:  Estimate_rounded by Theory
## F = 1.0378, num df = 85, denom df = 73, p-value = 0.8743
## alternative hypothesis: true ratio of variances is not equal to 1
## 95 percent confidence interval:
##  0.6621638 1.6142093
## sample estimates:
## ratio of variances 
##           1.037812
```

```
## 
##  Two Sample t-test
## 
## data:  Estimate_rounded by Theory
## t = 1.6608, df = 158, p-value = 0.09875
## alternative hypothesis: true difference in means is not equal to 0
## 95 percent confidence interval:
##  -0.0004970411  0.0057490838
## sample estimates:
## mean in group Not P-FIT     mean in group P-FIT 
##              0.01463953              0.01201351
```

##### c. Mean Thicknesses F-test, t-test

```
##      Theory   grp.mean
## 1 Not P-FIT 0.02219767
## 2     P-FIT 0.01704054
```

```
##      Theory     grp.sd
## 1 Not P-FIT 0.01718811
## 2     P-FIT 0.01381823
```

```
## 
##  F test to compare two variances
## 
## data:  Estimate_rounded by Theory
## F = 1.5472, num df = 85, denom df = 73, p-value = 0.05687
## alternative hypothesis: true ratio of variances is not equal to 1
## 95 percent confidence interval:
##  0.987184 2.406537
## sample estimates:
## ratio of variances 
##           1.547217
```

```
## 
##  Two Sample t-test
## 
## data:  Estimate_rounded by Theory
## t = 2.0688, df = 158, p-value = 0.04019
## alternative hypothesis: true difference in means is not equal to 0
## 95 percent confidence interval:
##  0.0002337114 0.0100805564
## sample estimates:
## mean in group Not P-FIT     mean in group P-FIT 
##              0.02219767              0.01704054
```

##### d. Plot of Distribution

#B) Interpretation P-FIT cortical volumes, thicknesses, and surface areas did not have larger associations with the g-factor than non-P-FIT regions with and without adjusting for global brain size (File S4). Non-P-FIT mean thicknesses (M = 0.02, SD = 0.02) had larger associations with the g-factor than P-FIT mean thicknesses (M = 0.01, SD = 0.01, t(143.49) = 4.84, p < 0.001) without adjusting for global brain size. Non-P-FIT mean thicknesses (M = 0.02, SD = 0.02) and volumes (M = 0.02, SD = 0.01) had larger associations with the g-factor than P-FIT mean thicknesses (M = 0.02, SD = 0.01, t(158) = 2.07, p < 0.05) and volumes (M = 0.01, SD = 0.01, t(157.7) = 3.19, p < 0.01) when adjusting for global brain size.
